# Influence of genetic interactions on polygenic prediction

**DOI:** 10.1101/667162

**Authors:** Zhijun Dai, Nanye Long, Wen Huang

## Abstract

Prediction of phenotypes from genotypes is an important objective to fulfill the promises of genomics, precision medicine and agriculture. Although it’s now possible to account for the majority of genetic variation through model fitting, prediction of phenotypes remains a challenge, especially across populations that have diverged in the past. In this study, we designed simulation experiments to specifically investigate the role of genetic interactions in failure of polygenic prediction. We found that non-additive genetic interactions can significantly reduce the accuracy of polygenic prediction. Our study demonstrated the importance of considering genetic interactions in genetic prediction.

## Introduction

The problem of “missing heritability” has attracted much attention and controversy in quantitative genetics (Manolio et al., 2009), yet its definition remains ambiguous in the literature. A widely used definition is that genetic associations identified in large-scale genome-wide association studies (GWAS) cannot fully account for heritability estimates (e.g. from twin studies) in the sense that the model fitting can only capture a fraction of the total variance. As sample sizes for GWAS increase from thousands to hundreds of thousands, and advanced statistical methods are developed to fit all DNA variants in the model simultaneously, including those not significantly associated with the trait, the variance that can be explained by DNA variants also increase. For example, adult human height is a classical quantitative trait with a narrow sense heritability (*h*^*2*^) of approximately 0.8 based on twin studies (Silventoinen et al., 2003). However, early GWAS studies identified common variants explaining only a total of 2-4% phenotypic variance (Gudbjartsson et al., 2008; Lettre et al., 2008; Weedon et al., 2008) with sample sizes in the order of 20,000. In 2010, a landmark study increased this proportion to about 45% by fitting ~300,000 SNP markers in the model for ~4,000 individuals with the covariance among individuals determined by genome-wide SNP similarity (Yang et al., 2010). Importantly, applying the same idea, the most recent study using whole genome sequences of ~20,000 individuals in the TOPMed almost entirely closed the gap between the genomic heritability and the presumed heritability (Wainschtein et al., 2019). The progress has been remarkable and it can be cautiously expected that the combination of large sample size and full genome sequences may finally capture all heritability. Perhaps more importantly, it also suggests that the failure to explain all heritability in early GWAS was due largely to low statistical power and incomplete variant coverage thus those with smaller effects, lower minor allele frequencies, and non-SNP variants were missed from the model fitting.

A second, more implicit but more practical definition of missing heritability, is that the prediction accuracy of quantitative phenotypes based on genotypes (polygenic scores) is far less than the heritability of the trait. A perfect genetic model with precise effects and model specification should be able to predict unobserved phenotypes with an accuracy (measured by *r*^*2*^) equal to the heritability. But that’s not always the case. For example, a large GWAS on adult human height with almost 200,000 individuals identified over 180 loci, capturing only ~10% of the phenotypic variation (Lango Allen et al., 2010). This proportion of variance was measured based on “leave-one-out” out-of-sample prediction (International Schizophrenia Consortium et al., 2009), *i.e.*, the effects of the genetic loci were estimated in one subset of the sample and polygenic scores (genetic effects summed over all significant loci) was computed to predict phenotypes in another subset. The partition between the subsets conveniently followed sample origin from different European countries (Lango Allen et al., 2010). In contrast to the mixed model genomic heritability approach, this method of estimating explained heritability was more akin to genomic prediction widely used in animal and plant breeding (Meuwissen et al., 2001; VanRaden, 2008), in which effects of genetic markers across the whole genome, regardless of their statistical significance, are summed to compute genetic prediction.

Both the genomic heritability and the prediction accuracy of polygenic scores (or polygenic breeding values) take the form of variance proportion, but have vastly different properties. One of the most contrasting differences is that prediction accuracy can be small even when the genomic heritability is large (Makowsky et al., 2011). Our discussion on the two definitions of missing heritability above is a clear example of this distinction. The implications of this distinction are profound. Most notably, even if there was no missing heritability based on genomic heritability, the utility of polygenic score would be very limited if prediction accuracy is low.

Recently, there has been renewed interest in the application of polygenic score (International Schizophrenia Consortium et al., 2009) with the advent of large public data sets such as the UK biobank (e.g. Khera et al., 2018). In particular, many studies have observed poor prediction by polygenic scores across different ancestry groups (Martin et al., 2019) or even within an ancestry group but with variable characteristics (Mostafavi et al., 2019). In fact, earlier studies with smaller sample sizes observed similar patterns, but were interpreted as missing heritability (Lango Allen et al., 2010; Makowsky et al., 2011). In animal breeding, similar observations have also been made. Although genomic prediction works exceedingly well within a breed, cross-breed prediction generally fails (Hayes et al., 2009). The explanation is obvious, genetic effects are context dependent and heterogeneous between groups. Variable linkage disequilibrium (LD) patterns, environments, and other factors can all contribute to the variable genetic effects, manifesting as variable accuracy of polygenic prediction.

Genetic interactions are pervasive, and an important type of context dependent effects (Mackay, 2014; Mackay and Moore, 2014). It has been previously shown that the presence of genetic interactions does not have a strong effect on genomic heritability (Hill et al., 2008; Huang and Mackay, 2016), therefore the magnitude of genomic heritability offers no indication of the genetic architecture. However, genetic interactions may influence genomic prediction accuracy and models taking into account the complexity improves prediction (Morgante et al., 2018). This clearly suggests that the simplification of genetic architecture to the additive infinitesimal model when the true model is not, although convenient and no comparable alternatives exist, can be risky. In this study, we specifically investigate the influence of genetic interactions on prediction of polygenic scores, with an emphasis on polygenic prediction across diverged populations.

## Results

### Experimental design

Because it’s not yet possible to unambiguously know the true genetic architecture of a quantitative trait, all experiments in this study were performed using simulated data instead of real data. This allows us to specifically ask simple questions while eliminating influence from other factors. We simulated a sample of 75,000 diploid individuals from three ancestry groups, where population A and B diverged 1,000 generations ago and their ancestors diverged from population C an additional 1,000 generations ago (Figure 1a). This specification is qualitatively similar to the global human population history where the ancestral population that went out of Africa were further split into multiple populations.

**Figure 1.**
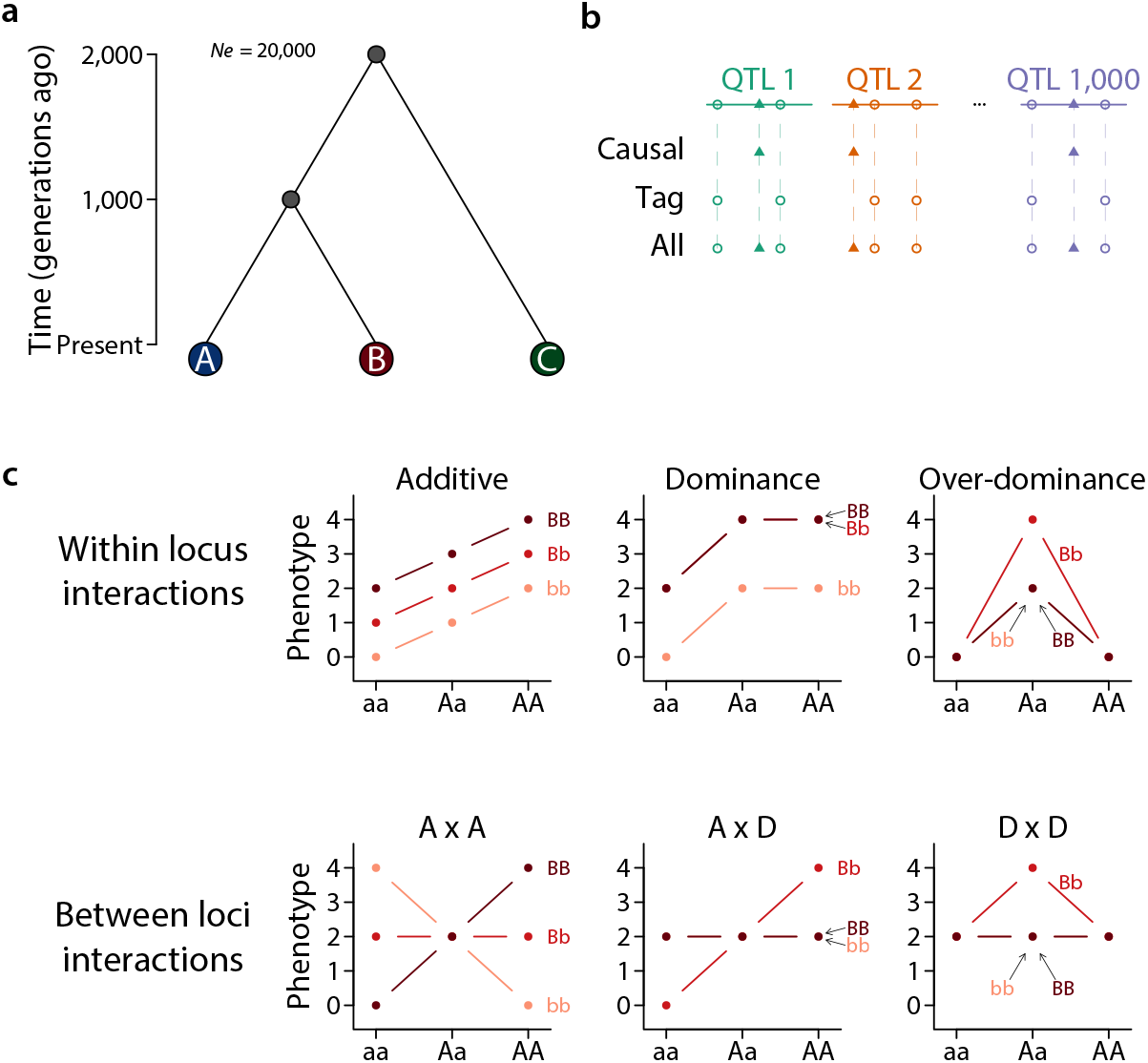
Simulation of genome sequences, population structure, and genetic architecture. **(a)** Three populations (A, B, C) were simulated with an effective population size of 20,000 each. A and B diverged 1,000 generations before present and A and C diverged 2,000 generations ago. **(b)** 1,000 independently inherited chromosomes were simulated, each containing one QTL. Three sets of variants were considered, including “causal”, “tag”, and “all” as illustrated. **(c)** Six different genetic architecture were simulated, each illustrated by one of the panels.

We considered three possible variant sets (Figure 1b), 1) causal: all and only causal variants; 2) tag: all variants except causal variants; and 3) all: all variants including causal variants. These represent three simplified scenarios 1) a best case scenario where causal variants have been identified, 2) a realistic scenario where causal variants are tagged by genotyped variants, and 3) an achievable scenario in the near future with whole genome sequences. We did not consider variants that were rare (MAF < 0.01) in all three populations as they led to gross overestimation of genomic heritability approaching one, similar to findings in a simulation study using real genotypes (Evans et al., 2018). The three variant sets were used to compute genomic heritability and perform polygenic prediction. When performing polygenic prediction, we did not select variants based on association tests. This choice was based on the consideration that selection of markers introduced another variable in the experiment to complicate the design and interpretation. Instead, we draw from the distinction between causal and all variants to represent the extreme scenarios where a perfect selection or no selection was performed.

We simulated a quantitative trait controlled by 1,000 independently inherited QTLs (Figure 1b) of broad sense heritability *H*^*2*^ = 0.8 but different types of genetic architecture. When the genetic architecture is strictly additive, the narrow sense heritability *h*^*2*^ = *H*^*2*^ = 0.8, whereas in other cases *h*^*2*^ < 0.8. Six simple models of genetic architecture were simulated, including additive, dominance, over-dominance, and pairwise additive by additive (A x A), additive by dominance (A x D), and dominance by dominance (D x D) (Figure 1c). No higher order interaction was simulated and effects across loci or across pairs were additive.

### Genomic heritability misses little heritability

We first recapitulated a result that has been consistently shown (Hill et al., 2008; Huang and Mackay, 2016). We fitted a linear mixed model in each of the three populations or combined samples using GREML implemented in the GCTA (Yang et al., 2011) using 20,000 individuals. We found that *h*_*g*_^*2*^ were uniformly high when the genetic architecture was additive, dominance, or additive by additive, accounting for nearly all heritability (Figure 2, Figure S1). Whether or not the variant sets included casual variants appeared to have little effects on *h*_*g*_^*2*^; variant sets excluding causal variants performed as well as causal variants only and there was a slight tendency of upward bias (Figure 2). Similar results were obtained regardless of whether the samples were from a homogeneous population or a mixture of samples from two diverged populations (Figure S1). When the genetic architecture was entirely overdominance, additive by dominance, or dominance by dominance, *h*_*g*_^*2*^ was lower, but still consistently explained > 50% of the heritability (Figure 2, Figure S1). Taken together, these results suggest that as long as a large number of genome-wide markers were fitted, little heritability was missed, regardless of the genetic architecture. In other words, the magnitude of genomic heritability offers no discrimination of the underlying genetic architecture (Huang and Mackay, 2016).

**Figure 2.**
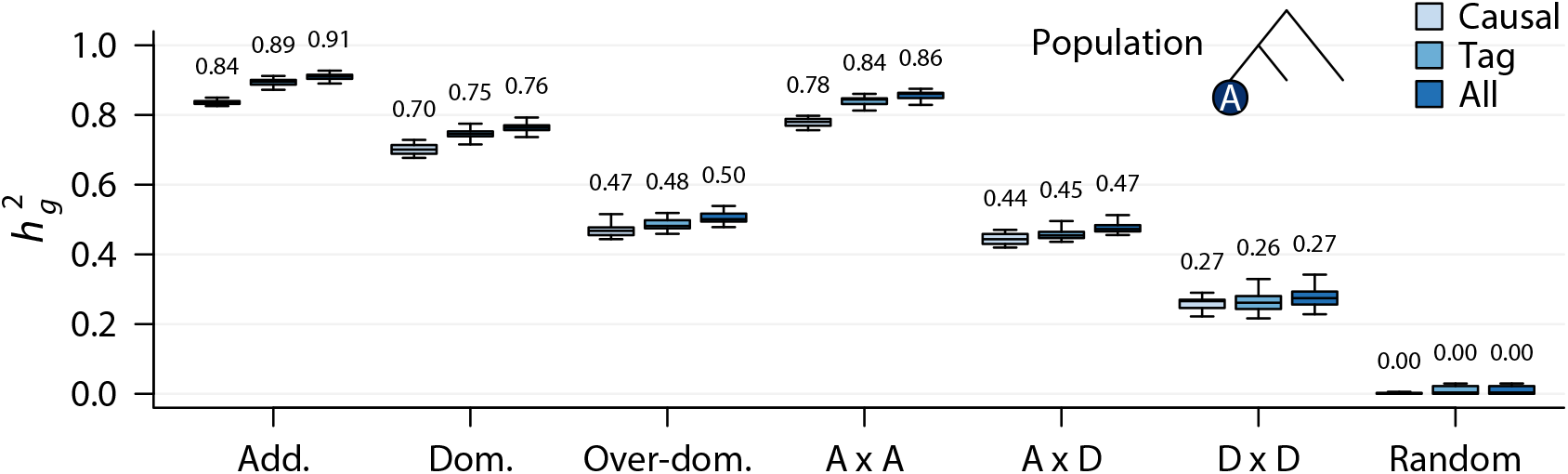
Genomic heritability in the simulated populations. Box plot (median indicated on top) showing the genomic heritability (*h*_*g*_^*2*^) estimated using GREML under different genetic architecture, where Add. = additive, Dom. = dominance, Over-dom. = over-dominance, A x A = additive by additive, A x D = additive by dominance, D x D = dominance by dominance, and random is a non-genetic model where the phenotypic variation was entirely due to random environmental variation. The population in which the genomic heritability was estimated was indicated in the top right corner. Genomic heritabilities in all populations were given in **Figure S1**.

### Accuracy of polygenic prediction with an additive genetic architecture

We then asked a simple question. If genome-wide variants are able to capture the majority of heritability, are they able to predict phenotypes accurately? This question directly addresses the distinction between the two definitions of missing heritability as we outlined in the introduction. If there is no missing heritability based on mixed model fitting, is there missing heritability in polygenic prediction? Many illuminating results could be obtained by comparing different scenarios of simulations (Figure S2).

We first considered the simplest and best scenario, in which the genetic architecture was fully additive, and all causal and only causal variants were known. In this case, the statistical model took the form of the true model and only model parameters needed to be estimated. We trained the model in one population (n = 20,000, training population) and computed polygenic scores of new individuals (n = 5,000, test population) either in the same population or a different population. To test the performance of cross-population prediction, we considered three possible relationships between the training and test populations, representing a gradient of divergence between training and test data (Figure 3a).

**Figure 3.**
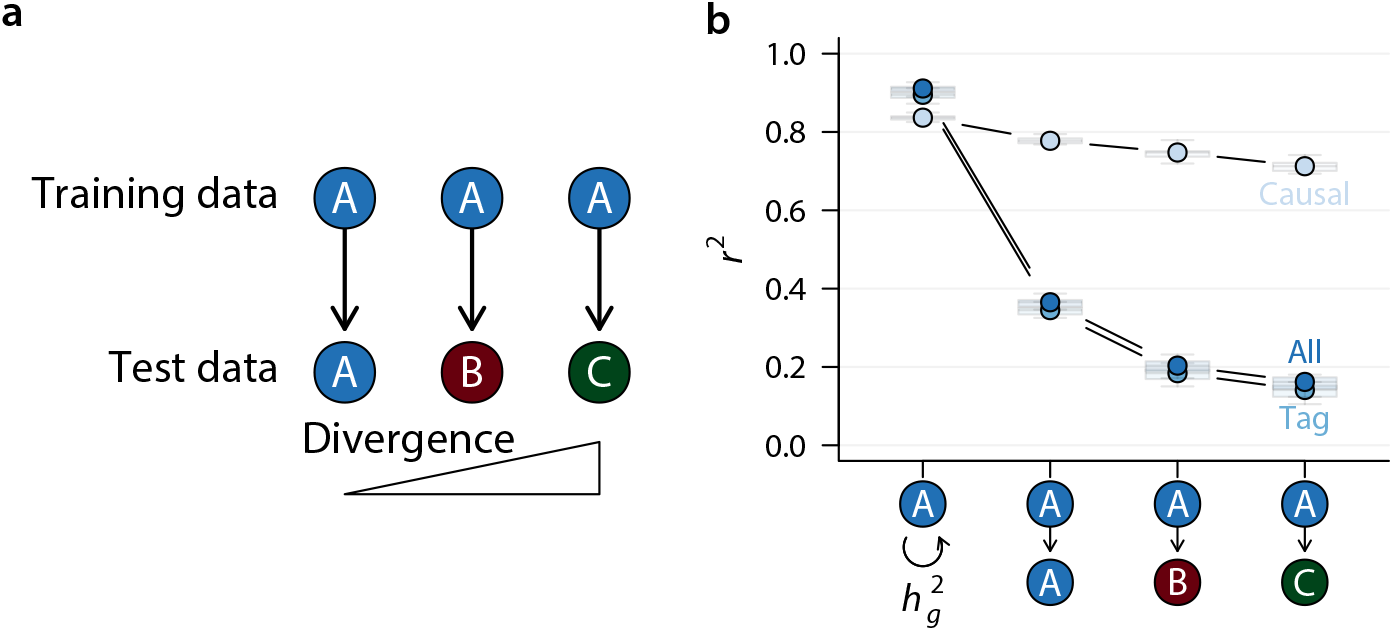
Polygenic prediction under additive genetic architecture. **(a)** Polygenic prediction was performed according to the diagram, where the model was trained in population A and tested in populations A, B, and C at increasing divergence. **(b)** Prediction accuracy was plotted according to the training – test population relationships. For comparison, genomic heritability was also plotted along side. Only the additive genetic architecture was considered in this plot.

As expected, the accuracy of polygenic prediction was very high in this best case scenario, approaching the true heritability (‘causal’ in Figure 3b). There was a small decline in accuracy when cross-population prediction was performed and the degree of population divergence negatively affected prediction accuracy. However, when non-causal variants were included to make predictions, accuracy plummeted from ~0.8 to ~0.4 (Figure 3b) even when training and test samples were from the same population. This was likely due to the inclusion of independent predictors whose number vastly exceeded that of the causal variants. As populations become more divergent, prediction accuracy further dropped, the rate of which was much more pronounced when tag or all variants were used. These results (in the cases of tag or all variant sets) largely agreed with the large body of empirical work that accuracy of polygenic prediction was substantially lower than genomic heritability and cross-population prediction was poor (Lango Allen et al., 2010; Makowsky et al., 2011; Martin et al., 2019).

One important lesson could be learned in this simple experiment. The facts that simply adding non-causal variants to the model drastically reduced prediction accuracy, and that the rate of decay in the accuracy of cross-population prediction was much greater in the presence of non-causal variants indicated that the agreement between model and true genetic architecture mattered. This is in sharp contrast to genomic heritability estimation, where including more variants generally improves model fit (compare (Yang et al., 2010) with (Wainschtein et al., 2019)).

### Accuracy of polygenic prediction in the presence of genetic interactions

We then tested the influence of genetic interactions on the accuracy of polygenic prediction, which fits an additive model. In a favorable condition when all causal variants were known (but not their effects or interactions) and prediction was performed within the same homogenous population, polygenic prediction accuracy was highly dependent on the genetic architecture (A -> A in Figure 4a). In general, prediction accuracy was higher for genetic architecture with higher *h*_*g*_^*2*^, such as additive, dominance, and additive by additive. In contrast, under overdominance, additive by dominance, and dominance by dominance genetic architecture, polygenic prediction performed substantially worse (A -> A in Figure 4a). When all variants were used, including non-causal ones, the prediction accuracies decreased dramatically and their dependency on genetic architecture appeared to be stronger (A -> A in Figure 4b).

**Figure 4.**
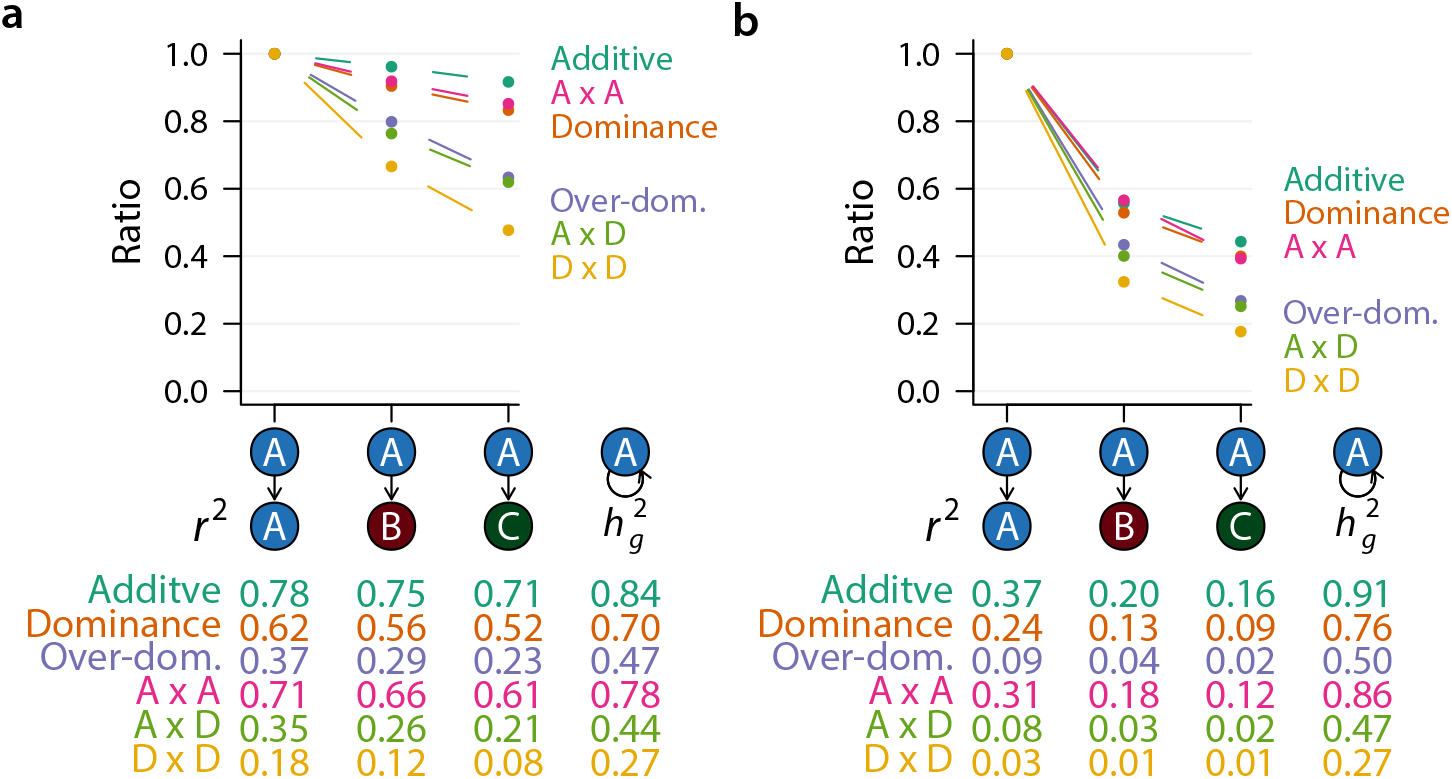
Polygenic prediction with different genetic architecture. **(a)** Polygenic prediction was performed using causal variants only for six different genetic architecture. The median prediction accuracy (*r*^*2*^) across 20 replicates in each scenario was listed below the graph, as well as genomic heritability (*h*_*g*_^*2*^). Each point on the graph represents a normalized median *r*^*2*^, dividing each prediction accuracy by its counterpart in the within population (A -> A) prediction. **(b)** Polygenic prediction with all variants. Data are presented the same way as in **(a)**. Data in these graphs were summarized from Figure S2.

We then asked how genetic interactions influence the rate of decay in prediction accuracies when the training and test populations diverge. We set the accuracy of within-population prediction as the baseline and compared cross-population prediction accuracies to this baseline. When all variants were used for polygenic prediction, the accuracy of cross-population prediction dropped to about 40-60% of the accuracy of within-population prediction, depending on genetic architecture (Figure 4b). Additive, additive by additive, and dominance genetic architecture, those with the highest *h*_*g*_^*2*^ and *r*^*2*^ retained the most prediction accuracy while over-dominance, additive by dominance, and dominance by dominance lost the most (Figure 4b). The more diverged the populations were, the more predictive ability of polygenic scores was lost (Figure 4b).

There are many reasons why polygenic prediction failed when test population diverged from training population. In our simple simulation setting, genetic effects were the same across populations and were not sensitive to any non-genetic factors. The difference in the linkage disequilibrium structure between populations may in part explain the drop. Importantly, simulations allowed us to directly use causal variants for prediction, thus eliminating the influence of LD (Figure 4a). Remarkably, while the accuracy of cross-population prediction was lower for all genetic architecture, the rate of decay was much greater when the genetic architecture was over-dominance, additive by dominance, or dominance by dominance (Figure 4a, compare slopes of the different lines). These results clearly suggest that genetic interactions can not only cause cross-population polygenic prediction to fail, but also in a more severe manner compared to an additive genetic architecture.

## Discussion

We demonstrate in this study through simulations that genetic interactions can influence the accuracy of polygenic prediction. In particular, cross-population polygenic prediction performed worse than intra-population prediction in all cases. For traits controlled by genetic interactions, the cross-population decay in prediction accuracy was far greater. The results make intuitive sense. For a statistical model to predict new data accurately, two conditions must be met. First, the model specification must be correct or at least sufficiently accurate to capture variation in the data. Second, parameters in the model must be precise. When genetic interactions are present, the additive polygenic model clearly is not accurate.

Previous studies have mostly focused on improving parameter estimation, through increasing sample size and methodological improvement. For example, increasing sample size substantially increased accuracy of polygenic prediction of height within individuals of European ancestry (Lello et al., 2018). Inclusion of samples of different backgrounds in the training data also helped (Martin et al., 2019, Figure S2).

However, the complexity of the genetic architecture of a quantitative trait makes it nearly impossible to specify a model prior to modeling. As a consequence, the polygenic infinitesimal model or variants of it (Gianola et al., 2009) has been used as the default model. The infinitesimal model has been instrumental and allowed for many theoretical insights as well as applications to be developed. In particular, prediction of breeding values in animal and plant breeding relying on the infinitesimal model has been very successful (García-Ruiz et al., 2016). However, its limitations are also apparent. Cross-population and cross-breed polygenic prediction was low in accuracy (Hayes et al., 2009; Lango Allen et al., 2010; Martin et al., 2019). Although many factors may contribute to this limitation, our simulation results clearly indicated that genetic interactions unaccounted for was a major contributor. Indeed, if the correct genetic model could be specified, cross-population prediction can achieve very high accuracy (Figure 5). There have been attempts to explicitly model non-additive genetic effects in the context of polygenic prediction; some moderate improvement was observed (Varona et al., 2018). However, these studies modeled non-additive effects using genome-wide markers, which added a large number of independent predictors as noise to the model and may negatively impact the performance.

**Figure 5.**
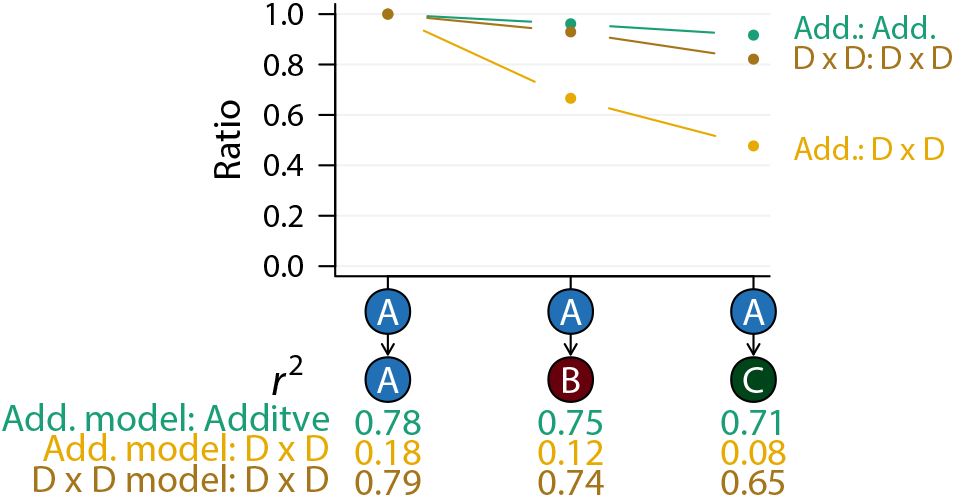
Match between model and true genetic architecture improves polygenic prediction. Two genetic architecture were considered, the additive and D x D. The prediction was performed with either an additive model (Add. model) as implemented in GREML or a D x D model in which the correct genetic model was presumed to be known and fitted. Only causal variants were used in these analyses.

We did not analyze existing large data sets, some of which contained subjects from multiple ancestries. Previous work with real data has consistently shown that cross-population polygenic prediction generally fails (Martin et al., 2019). However, it is difficult to disentangle the different factors that may contribute to effect heterogeneity and the failure of prediction in real data sets. Using simulations, we can focus on specific questions and our results clearly indicated a contribution of genetic interactions to the failure of cross-population polygenic prediction. While the additive infinitesimal model is the most sensible model when no other information is available, our study suggests that the development in the field should be expanded to include efforts to more explicitly model genetic interactions. Although it is challenging, recent advances in modeling (Boyle et al., 2017; Liu et al., 2019) and genomic assays informing regulatory networks (Gerstein et al., 2012) may finally offer new ways to develop biologically sensible models.

## Methods

### Population simulation

We used the coalescent simulator MaCS (Chen et al., 2008) to simulate genome sequences of 75,000 individuals, with 25,000 in each of the populations, according to the demographic history in Figure 1a. We simulated 1,000 independently inherited chromosomes of 100,000 base pairs in size and set mutation rate as 1.25 × 10^−8^ per bp and recombination as 1.25 × 10^−8^ per bp. Effective population size was set to 20,000. The MaCS command for one chromosome was `“macs 150000 100000 -s "$random_seed" -i 1 -h 1000 -t 0.001 -r 0.001 -I 3 50000 50000 50000 0 -ej 0.0125 3 2 -ej 0.025 2 1”`. This simulation was performed once but the partition between samples were repeated 20 time, which were summarized as box plots in figures.

### Simulation of quantitative phenotypes

We simulated quantitative phenotypes according to the genetic architecture depicted in Figure 1c. For each of the three possible genotypes for a biallelic locus with alleles A and a, we used the additive coding aa = −1, Aa = 0, and AA = 1 and the dominance coding aa = 0, Aa = 1, AA = 0 to code genotypes. The simulation of phenotypes consisted of two steps. In the first step, the corresponding genotype coding for an individual or multiplication of genotype codings (in the case of between-loci interactions) were multiplied by a genetic effect randomly drawn from the standard normal distribution and summed over all loci or all pairs of loci to obtain the genetic values. In the second step, an environmental effect was added by drawing from a normal distribution with a computed variance such that the broad sense heritability *H*^*2*^ = 0.8. We performed this simulation in each of the 20 random partitions of populations and independently sampled causal variants and genetic effects.

### Fitting GREML

We fitted the GREML model using GCTA (Yang et al., 2011) with 20,000 individuals from each of the A, B, and C populations and A + B and A + C. The GREML partitioned phenotypic variance into a genomic (*σ*^*2*^_*g*_) and an environmental component (*σ*^*2*^_*e*_). Genomic heritability was computed as *h*^*2*^_*g*_= *σ*^*2*^_*g*_/(*σ*^*2*^_*g*_ + *σ*^*2*^_*e*_).

### Polygenic score prediction

The BLUP estimates of SNP effects were obtained using GCTA and provided to PLINK2 (https://www.cog-genomics.org/plink/2.0/credits) to compute a polygenic score in 5,000 new individuals either from the same population as the fitted model or from a different population. Prediction accuracy of polygenic score was computed as the *r*^*2*^ of correlating predicted polygenic scores and the simulated true phenotypes. In the case of prediction using causal variants with the correct dominance by dominance model (Figure 5), we constructed pseudo-variants using the relevant genotype coding (for D × D, double heterozygotes were coded as one genotype class and all others another) and ran GREML and polygenic score prediction the same way as an additive model.

---

**Figure S1.**
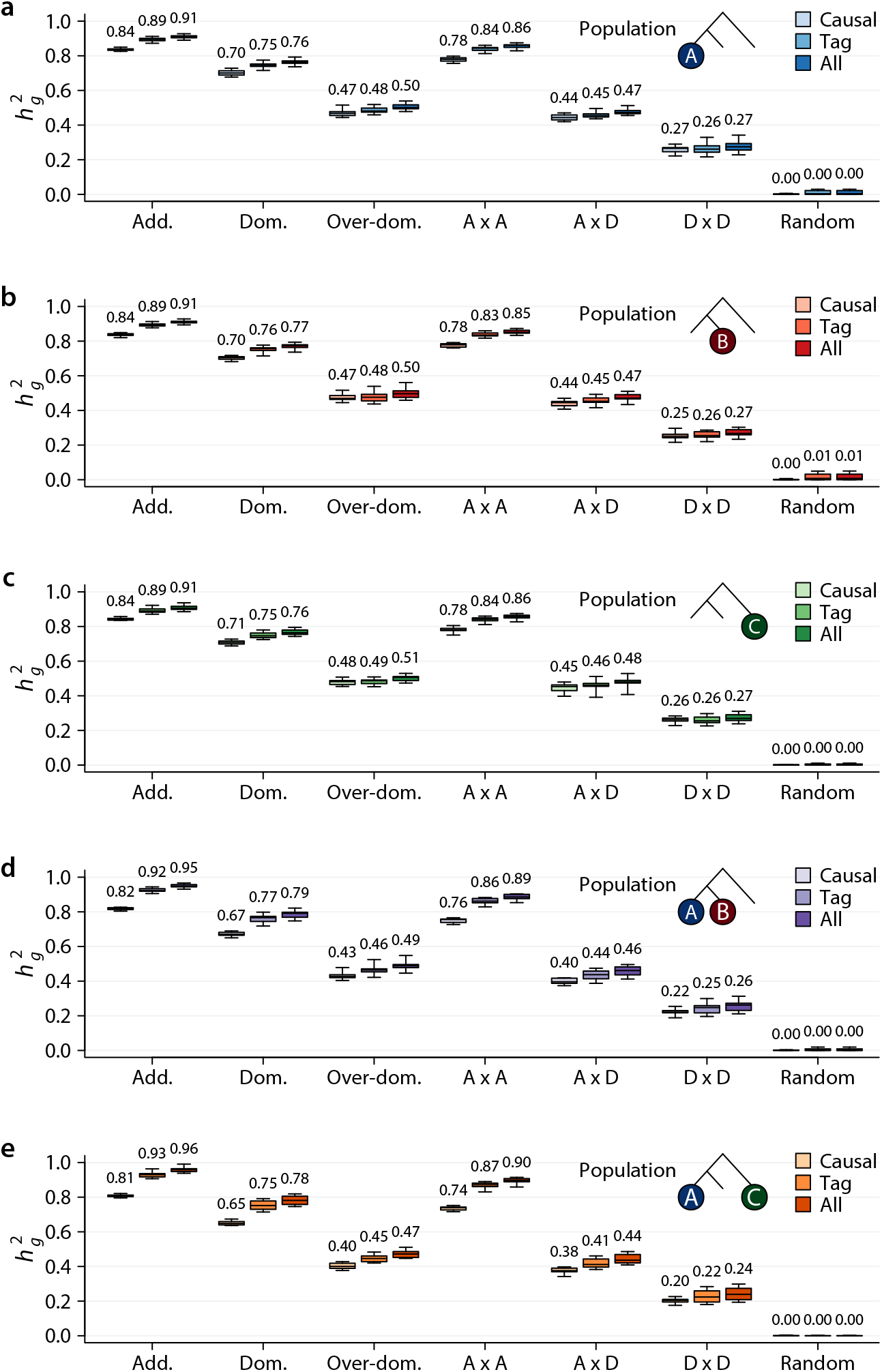
Genomic heritability in different simulated populations. Genomic heritability was plotted for different population samples. **(a)** population A; **(b)** population B; **(c)** population C; **(d)** population A + B; and **(e)** population A + C.

**Figure S2.**
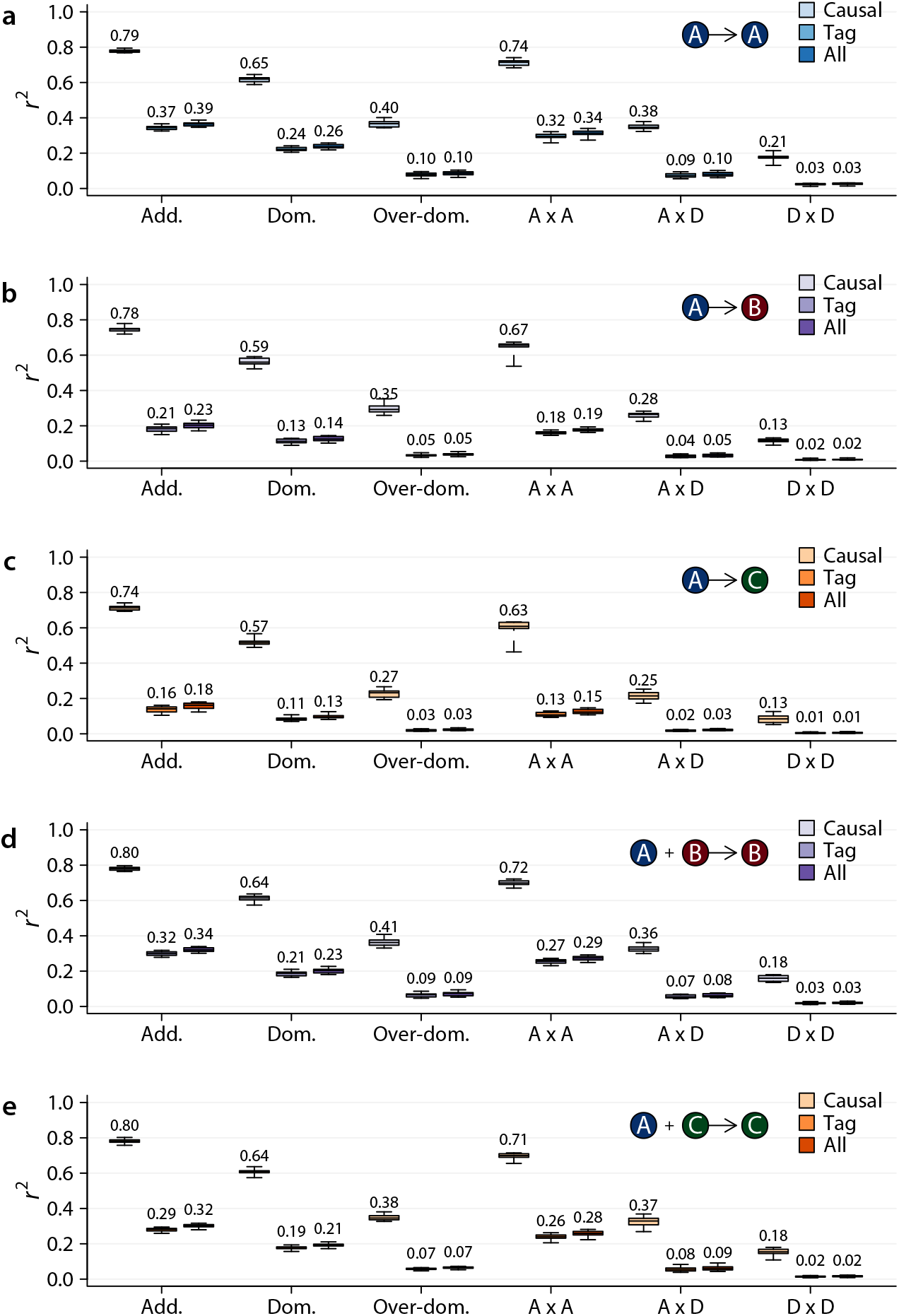
Polygenic prediction under different genetic architecture in different populations. Accuracies of polygenic prediction under different genetic architecture in **(a)** fit model in A, predict in A; **(b)** fit in A, predict in B; **(c)** fit in A, predict in C; **(d)** fit in A + B, predict in B; **(e**) fit in A + C, predict in C.

